# Hybrid capture RNA-seq defines temporal gene expression in *Rickettsia*

**DOI:** 10.64898/2026.01.04.697603

**Authors:** Allison T. Scott, Jon McGinn, Vincent L. Butty, Stuart S. Levine, Rebecca L. Lamason

## Abstract

Pathogenic *Rickettsia* species are obligate intracellular bacteria that must reside in a mammalian host or arthropod vector cell to survive. Although these bacteria transition between different intracellular environments during infection, they encode few putative transcription factors, and their gene regulatory networks are largely unknown. Because of their inextricable relationship with eukaryotic cells, transcriptional profiling of the pathogen is complicated by the abundance of contaminating host RNA, especially in infection conditions or stages where the bacterial burden is inherently low. Here, we employ a hybrid capture technique (PatH-Cap) to improve library preparation by enriching bacterial transcripts while depleting host and rRNA molecules. Using PatH-Cap, we explored transcriptional changes throughout the first 24 hours of infection, including infection initiation – an infection stage which has been difficult to profile with standard library preparation methods. We then clustered genes based on their temporal trends, revealing cohorts of genes whose expression is up- or downregulated at different stages of infection. We also highlighted the diverse temporal expression trends of genes with known roles in growth and pathogenesis, including translation and cell division genes, secreted effectors, and secretion system components. Lastly, we identified 310 antisense RNA molecules, many of which also showed strong temporal trends. This work demonstrates that sensitive transcriptional profiling approaches like PatH-Cap hold great promise for dissecting gene expression networks driving infection in intracellular pathogens that have historically posed significant technical challenges.

**IMPORTANCE:** When investigating poorly annotated genomes, such as those in obligate intracellular bacteria, transcriptional analyses can reveal gene sets active under specific conditions and form the foundation for future targeted approaches. However, such systems-level analyses of dynamic gene expression changes during infection with *Rickettsia* species have been missing due to the limitations of standard RNA-seq library preparations. Here, we adapted the Pathogen Hybrid Capture (PatH-Cap) method for the first time to any *Rickettsia* species. We leveraged this wealth of RNA-sequencing information to compare temporal trends between genes and investigate aspects of *Rickettsia parkeri* transcription regulation, such as predicting operon structure and identifying putative antisense RNA transcripts. This work establishes the most comprehensive analysis of temporal rickettsia gene expression to date, providing an important foundation for further analysis. Future work can apply the methods described here to investigate gene expression changes across different genetic or environmental perturbations, cellular contexts, or disease models.

## INTRODUCTION

Spotted fever group *Rickettsia* spp. are tick-borne obligate intracellular bacteria, many of which cause life-threatening vascular diseases in humans (1). Due to their inextricable interaction with host and vector cells, *Rickettsia* spp. have evolved a litany of mechanisms to manipulate eukaryotic cells and evade immune responses throughout their complex lifecycle. These lifecycle stages include escaping from host vacuoles, two temporally distinct forms of actin-based motility, replication, and spread between host cells. Research over decades has explored the mechanistic underpinnings of these host-microbe interaction (1), leading to the identification of secreted and surface proteins that are leveraged by the bacterium to hijack host processes. However, the ways in which the bacterium senses environmental cues within the host and execute specific gene expression programs to transition through these lifecycle stages remains unknown (2).

Reductive evolution has led to extremely small genomes among *Rickettsia* spp. resulting in genomes of approx. 1.3 million base pairs with an average of only 1200 genes. A large fraction of annotated genes within these streamlined genomes encode hypothetical proteins, which lack functional annotations and are often restricted to the *Rickettsia* genus. Even when exploring well-studied biological pathways, genomic inversion and deletion events have led to a high number of pseudogenes (3), and many core metabolic (4), replicative, and DNA-maintenance (5) processes seem to be lacking common components. *Rickettsia* spp. also encode very few transcriptional regulators (2), raising the question of how gene expression programs across host species or throughout the infectious cycle are mediated, and suggesting that alternative methods of gene regulation such as regulatory RNAs (sRNA), may play a significant role (6–9). Prior research has illuminated the unique biology *Rickettsia* spp. employ to manipulate their host, survive in the intracellular space, and respond to their environment, but many open questions remain (10, 11). Transcriptional profiling can add to this body of work by providing critical insight into the expression patterns of novel genes, which can bolster targeted analysis or inference of gene function from bioinformatic predictions, but most studies of rickettsial transcriptomics are limited in scope, due to technical difficulties.

RNA sequencing of intracellular pathogens is complicated by the abundance of host RNA molecules present in total RNA extractions of infected cells. One calculation estimates as little as 0.25% of total RNA from infected cells originates from bacterial sRNA and mRNA (12). Previous reports investigating rickettsial transcription have increased this percentage using a combination of techniques, such as massively increasing the infectious burdens and read depth, removing eukaryotic mRNA by polyA depletion, and depleting host and bacterial rRNA. Even with these multi-step enrichment processes, however, the percent of total reads mapping to the rickettsial genome often remains low and is highly dependent on initial infectious burden, infection duration, and the infection system (2, 6, 7, 13–16). Although these studies were successful in broadly profiling bacterial transcription in mammalian or arthropod cells and validating the presence of sRNAs, we lack gene expression information during early stages of infection when bacterial burdens are low and have no information on how gene expression changes during the infectious cycle. Recently, studies using infection models of *Plasmodium vivax* (17), *Pseudomonas aeruginosa* (18), and *Mycobacterium tuberculosis* (18, 19) have considered similar challenges and consequently developed a positive-selection method called PatH-Cap (Pathogen-Hybrid Capture) for more robust transcriptional profiling. This technique relies on bespoke probe libraries that hybridize with and enrich pathogen transcripts while simultaneously depleting host, rRNA, and tRNA transcripts (**Fig. 1A**). PatH-Cap was used successfully to profile the transcription of 1–3 bacteria per host cell (18), demonstrating the power of this approach for low input samples.

**Fig. 1.**
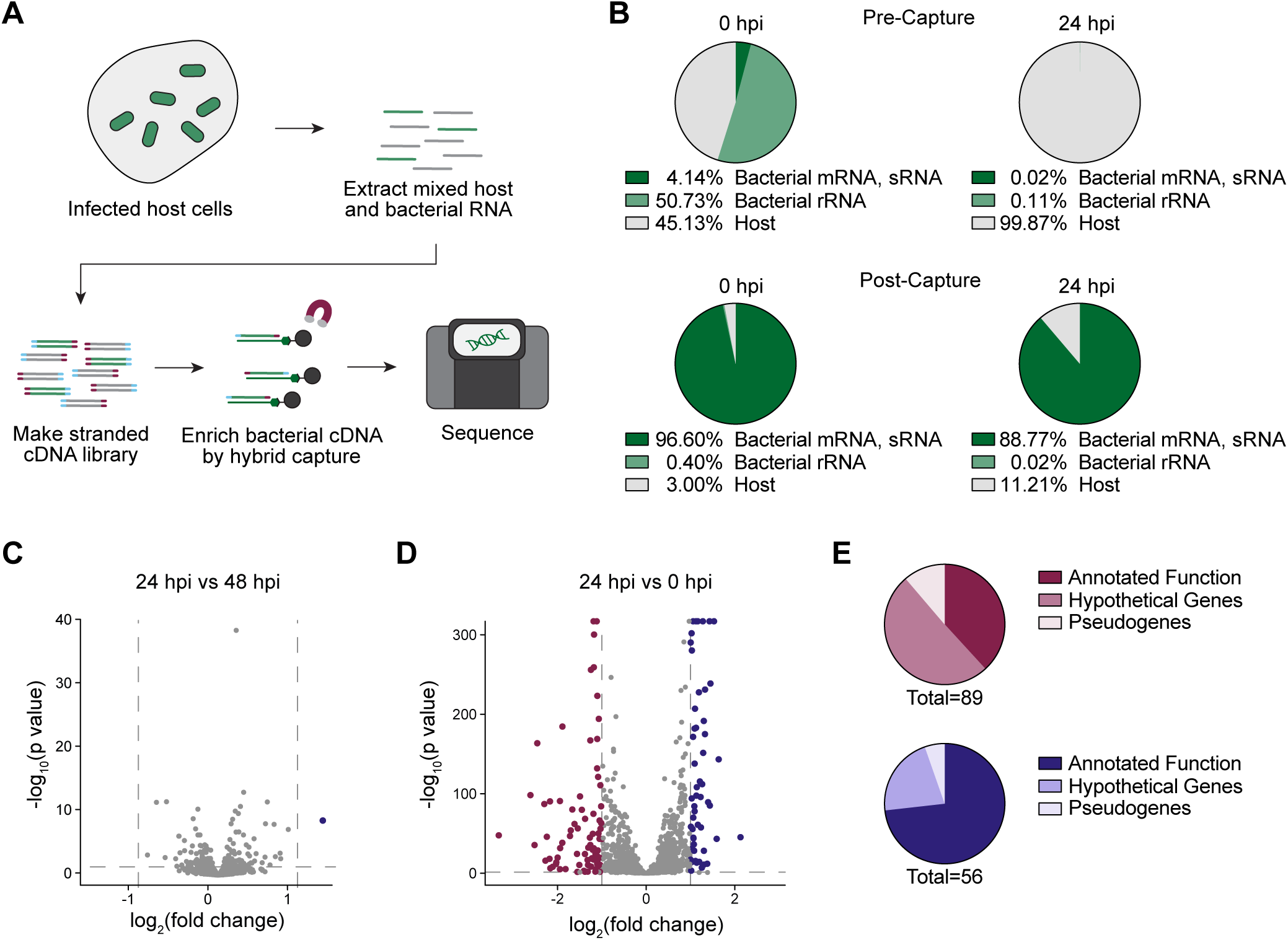
PatH-Cap enriches for bacterial transcripts from *R. parkeri*-infected host cells. (**A**) Diagram of PatH-Cap adapted for *R. parkeri*. Bulk RNA from infected cells was fragmented, barcoded, and reverse transcribed into a double stranded cDNA library. Biotinylated probes that tile the *R. parkeri* genome were hybridized to bacterial-derived cDNA, eluted, and sequenced. (**B**) Comparison of sequenced read content before and after PatH-Cap. Percentages were averaged across 8 biological replicates. (**C**) Differentially expressed genes comparing samples collected at 24 hpi and 48 hpi. (**D**) Differentially expressed genes comparing samples collected at 24 hpi and 0 hpi. Dotted lines indicate genes down-regulated (red) or up-regulated (blue) at 24 hpi with cutoffs of log_2_(FC) > 1 or < −1 and p value < 0.05. (**E**) Differentially expressed genes at 24 hpi relative to 0 hpi divided by annotated function, hypothetical genes, and pseudogenes. Red chart represents down-regulated genes and blue chart represents up-regulated genes. (C–E) Data are from 8 biological replicates.

In this study, we apply PatH-Cap to temporally profile the infectious life cycle of *Rickettsia parkeri*, a model SFG *Rickettsia* species, in mammalian host cells. We show that PatH-Cap allows robust detection of bacterial transcriptional changes even at a low MOI and throughout the early stages of infection. With this improved capacity, our study reveals for the first time networks of genes that display temporal signatures, highlighting their potential for stage- or context-dependent functions. Moreover, we expand the identity of *cis*-encoded antisense RNAs whose own temporal signatures suggest various modes of gene expression regulation.

## RESULTS

### Establishing PatH-Cap for *R. parkeri*

PatH-Cap was reported to dramatically enrich the transcripts of several pathogens under physiologically relevant infection conditions (18), suggesting it could be applied to profile the rickettsial transcriptome. Therefore, we designed an *R. parkeri*-specific hybrid capture library with 10,746 non-overlapping 120-nucleotide probes tiling one strand of the *R. parkeri* genome, including noncoding regions and excluding rRNA and tRNA genes. To quantify PatH-Cap enrichment of *R. parkeri* transcripts, we infected human epithelial cells (A549) with wild-type *R. parkeri*, since key bacterial life cycle events have been readily established in this cell culture system (20, 21). Total RNA was collected at 24 hours post infection (hpi) and from purified input bacteria (0 hpi) and used to generate a double stranded cDNA library, which preserved the strand information of the original RNA molecule. We used our biotinylated *R. parkeri* probe set to enrich for bacterium-derived transcripts and sequenced the pre- and post-capture libraries (**Fig. 1A** and **B**). While our pre-capture samples had the expected high levels of host contamination, our 0 and 24 hpi post-capture samples had 96.60% and 88.77% of reads, respectively, mapping to bacterial mRNA and sRNA (**Fig. 1B**). With this improved library preparation method, we obtained an average read depth of 536 reads per base with only an average sequencing depth of 5.2 million reads per replicate for our 24 hpi samples. These data demonstrate that we can produce robust read depth with a fraction of the sequencing depth as previous work. We also compared the consistency of this enrichment between two independent experiments and observed high correlation between experiments (R^2^ = 0.92 at 0 hpi and R^2^ = 0.94 at 24 hpi, **Fig. S1A** and **B**). Unlike other bacterial systems where PatH-Cap has been applied, the *R. parkeri* genome is highly AT rich with a GC content of only 32.5% (22). Importantly, we did not see any significant correlation between GC content and expression at the gene level (R^2^ = 0.014, **Fig. S1C**). Thus, PatH-Cap functions in bacteria with diverse genome composition and successfully enriches for *R. parkeri* transcripts in samples dominated by host transcript contamination.

### PatH-Cap enables fine time-course transcriptional profiling

We first used this pipeline to identify differentially expressed genes at 0, 24, and 48 hpi. We identified only one differentially expressed gene (*rnpB*) when comparing the 24- and 48-hour time points (**Fig. 1C**), in line with previous studies that have investigated transcriptional patterns during late-stage infection (13). Intriguingly, however, there were many more differentially expressed genes when comparing the 0- and 24-hour time points (**Fig. 1D**, **Table S1**), including many hypothetical and pseudogenes (**Fig. 1E**), suggesting *R. parkeri* may undergo dynamic transcriptional changes while establishing infection. These results prompted us to explore a finer-scale temporal analysis of *R. parkeri* gene expression by collecting RNA from A549 cells infected with wild-type *R. parkeri* at an MOI of 1 at 0, 1, 4, 8, 12, 16 and 24 hpi (**Fig. 2A**) – timepoints which capture distinct known stages of *R. parkeri* intracellular infection. Notably, we observed robust post-capture enrichment of bacterial mRNA reads even at 1 and 4 hpi, with 74.4% and 69.13% of reads, respectively, mapping to bacterial mRNA and sRNA. Together, these data indicate that PatH-Cap is compatible with early time points and as little as a single bacterium per host cell.

**Fig. 2.**
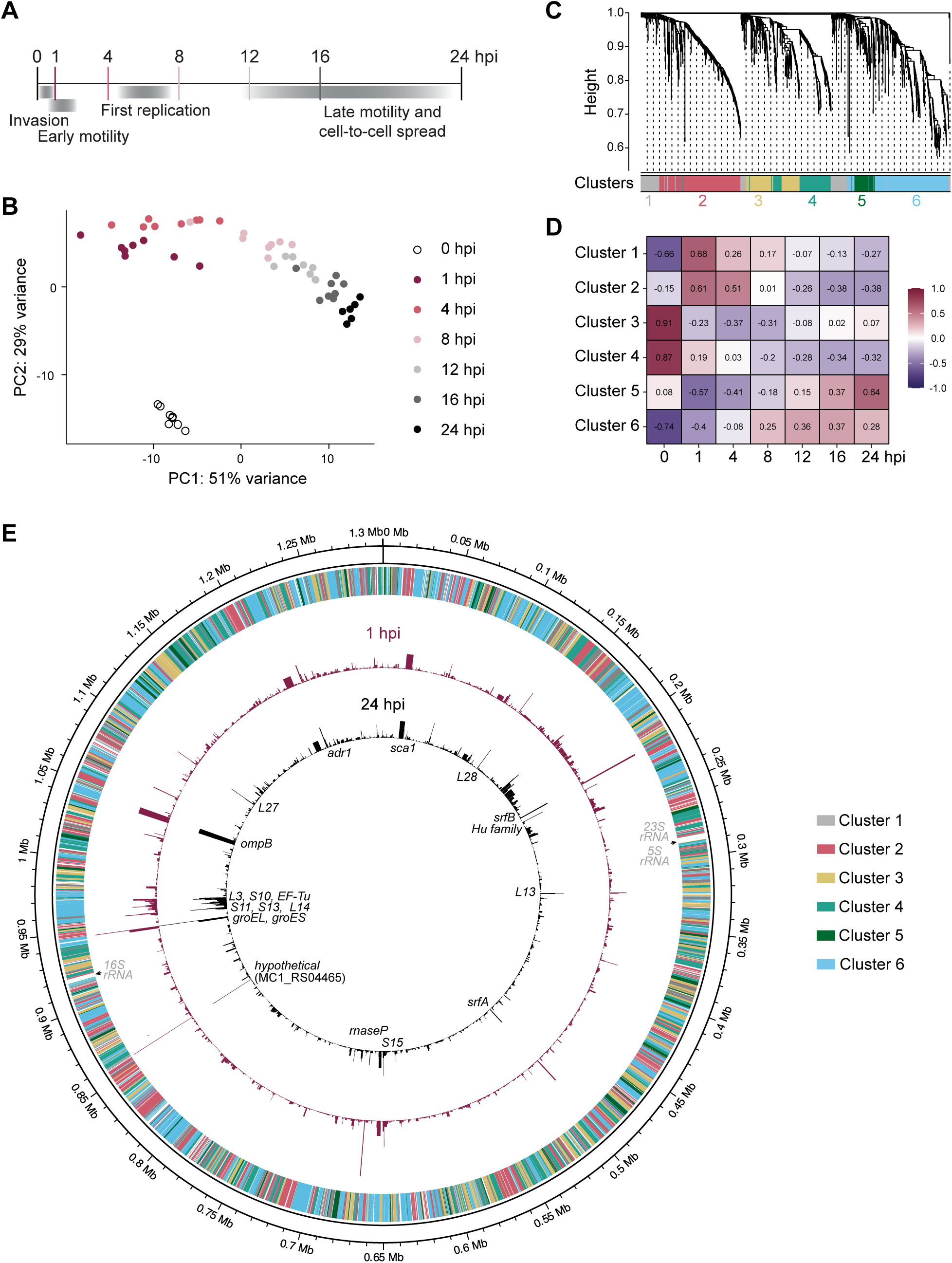
Time course RNA-seq profiles *R. parkeri* transcription during the first 24 hours of infection. (**A**) Diagram of sample collection time points and the corresponding *R. parkeri* life cycle stage. (**B**) Principal component analysis of time course samples. Data were regularized-logarithm (rlog) transformed using DESeq2 package in R prior to analysis. (**C**) Cluster dendrogram shows gene cluster assignments based on WGCNA. (**D**) Heatmap shows correlation between gene clusters and time points. Pearson correlation coefficient is reported for each cluster-time point pair. (**E**) Genome diagram shows each gene’s cluster assignment (outer ring), and average TPM at 1 hpi (red) or 24hpi (black). 20 genes with the highest expression are annotated in black, and the excluded rRNA genes are annotated in gray. Average TPM values are from 6–8 biological replicates per time point.

We next sought to evaluate the relationship between our samples using principal component analysis (PCA) (**Fig. 2B**). The 0 hpi samples clustered separately from the other time points, which is expected considering the different environment encountered by bacteria purified from host cells. In contrast, samples from 1–24 hpi clustered in temporal order, consistent with progressive changes in gene expression over time. We found that the 20 highest and lowest-expressed genes were similar across the 1 – 24 hpi samples (**Table S2**). Consistent with other studies (13, 14), the genes with the highest overall expression include genes encoding ribosomal proteins, outer membrane proteins OmpB and Sca1, and the chaperones GroEL and GroES. Additional noteworthy members include genes encoding SrfA and SrfB, which we previously isolated as novel secreted effectors (23), and the outer membrane protein Adr1, which may promote complement resistance (24). Most of the lowest expressed genes have not been functionally characterized, but several have distinct expression patterns in conditions different from our model system. For example, homologs of MC1_RS02615 and MC1_RS09865 were upregulated in tick cells relative to mammalian cells (14), and the MC1_RS06575 homolog was upregulated in response to low temperatures (2). Overall, PCA demonstrates *R. parkeri* undergoes temporal transcriptional changes that are not driven by fluctuations in genes with the highest and lowest expression, necessitating more global analysis of these trends.

### WGCNA facilitates clustering of temporally related rickettsial genes

We next turned to weighted gene co-expression network analysis (WGCNA) (25), which has been used in other works to cluster genes with highly similar transcriptional patterns (26, 27). We selected a soft power value to maximize our data’s fit with a scale free topology model (**Fig. S2A**) and decrease mean connectivity (**Fig. S2B**). Because the *R. parkeri* genome is small and contains many hypothetical genes, we set minimum cluster size and cut height values to avoid production of many smaller clusters dominated by genes of unknown function (**Fig. 2C**). Our analysis resulted in 6 different clusters of genes (**Fig. 2C**, **Table S3**). Each cluster is defined by a unique eigengene vector (25), which can be used to calculate the correlation between clusters and time points (**Fig. 2D**). Because we used a signed network model for WGCNA, clusters that are positively or negatively correlated with a particular time point contain genes that have increased or decreased expression at that time point, respectively. For example, genes in cluster 2 display peak expression during early infection (1–4 hpi) (**Fig. 2D**). Genes in clusters 3 and 4 have decreased expression over time, in contrast to genes in clusters 5 and 6, which generally show increased expression over time (**Fig. 2D**). Genes that lacked obvious temporal signatures in our analysis were grouped into cluster 1 (**Fig. 2C** and **D**, **Fig. S3**).

### Gene expression does not obviously correlate with genome position

In well-studied bacteria like *Escherichia coli* and *Bacillus subtilis*, genes are often spatially patterned in genomes (28). For example, genes near highly expressed genes (e.g., ribosome components) or the origin of replication may have higher expression relative to other regions (29). The *Rickettsia* genome, however, has undergone various recombination, acquisition, and reduction events that have scattered genes and sacrificed synteny (3, 30), leading us to ask whether a similar relationship between chromosomal position and temporal gene expression existed in *R. parkeri*. We first plotted the TPM for each gene at 1 hpi and 24 hpi (**Fig. 2E**) and noted that some highly expressed ribosomal and translation-associated genes are positioned together on the chromosome, but these loci are not proximal to the origin of replication (**Fig. 2E**). This positioning differs from *E. coli* where replication of the origin increases the gene dosage of these critical factors during their rapid 20 min replication cycle (28, 31). SFG *Rickettsia* spp. replicate over much longer time scales of 5-6 hours (21, 32), which may explain why positional control is not a conserved feature. We then plotted the cluster identity for all genes across the *R. parkeri* chromosome and found that the clusters were scattered (**Fig. 2E**), highlighting that these networks were not directly influenced by genomic position.

### Clusters contain differentially expressed genes with variable functional classifications

WGCNA groups genes with highly correlated expression patterns, even if the dynamic range of those changes is quite variable, requiring us to explore the extent of differential gene expression in each cluster. We calculated the log_2_(fold change) for each pairwise time point comparison and identified the maximum log_2_(FC) value, and found that most genes in these clusters exhibited at least a 50% change in gene expression over time (**Fig. 3A**). We then plotted the Z-score of genes in each cluster (**Fig. 3B**), which recapitulated the temporal trends determined by the cluster eigengenes above (**Fig. 2D**). Lastly, we defined gene ontology (GO) terms for functional enrichment analysis within each cluster using InterProScan (33). GO term assignment in *R. parkeri* is hindered by the abundant hypothetical genes that lack significant homology outside the genus, so we manually edited the predictions to ensure consistency with the literature. After assigning predicted GO terms (**Table S3**), 6.89% (103 genes) were classified as pseudogenes and 29.85% (466 genes) remained hypothetical. Among the genes with annotated functions, distinct GO terms were enriched in separate clusters, as indicated by fold enrichment values above 1 (**Fig. 3C, Table S4**). Only a few terms reached statistical significance, as discussed below, likely due to the small genome and high number of hypothetical genes.

**Fig. 3.**
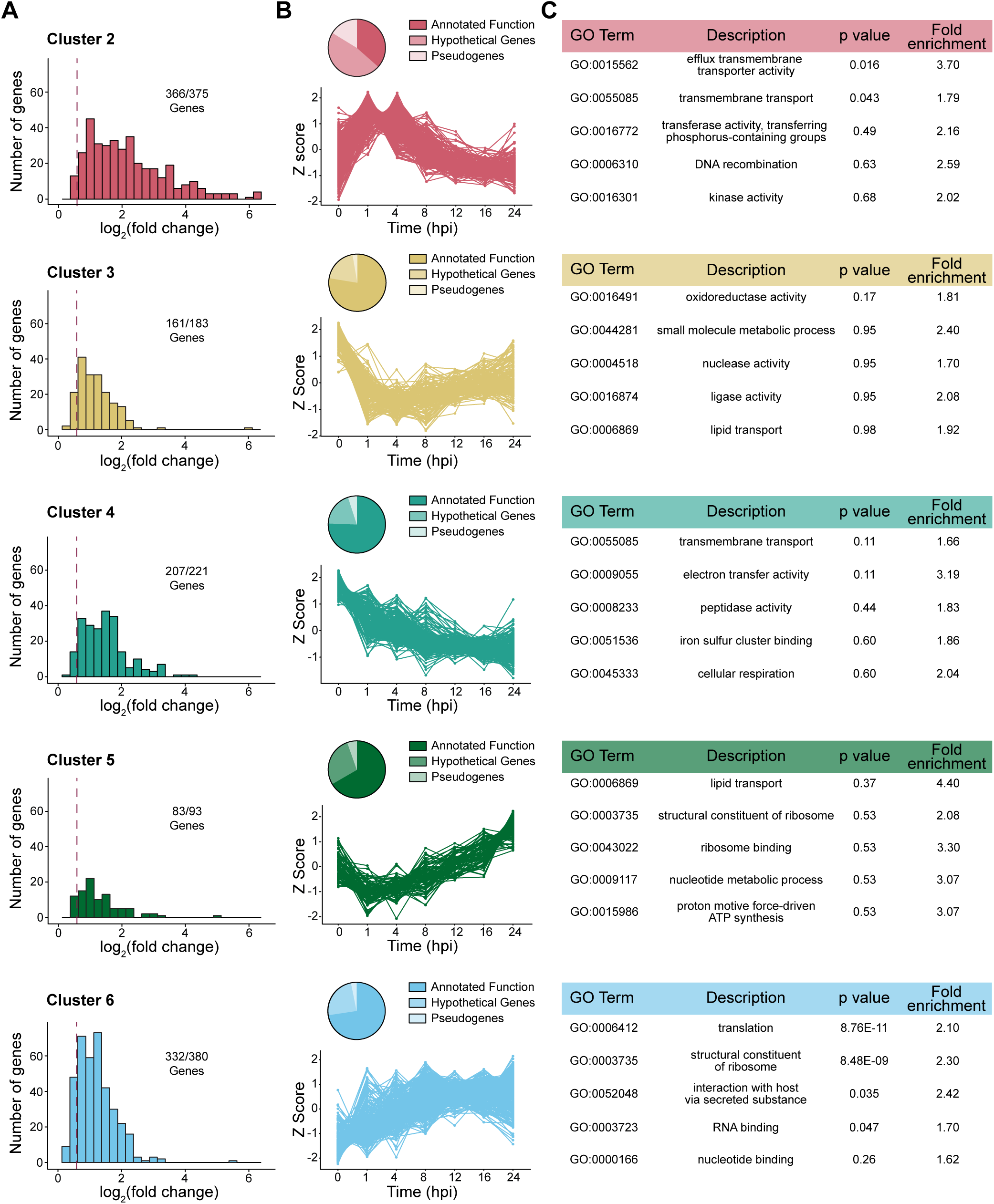
Gene clusters show distinct temporal trends and component genes. (**A**) Histogram log_2_(FC) for genes in each cluster. Maximum log_2_(FC) values were determined by computing DESeq2’s pairwise differential expression analysis for all possible time point combinations, identifying the log_2_(FC) value with the largest magnitude, and graphing the absolute value of the maximum log_2_(FC) for each gene. Dashed line represents log_2_(FC) = 0.58 and the number of genes per cluster above this threshold are reported. (**B**) Pie charts show the proportion of genes within a cluster annotated as genes of annotated function, hypothetical genes, and pseudogenes. Line graph shows Z-score over time of genes in each cluster. (**C**) Top 5 enriched GO terms within each cluster, reported regardless of statistical significance. GO term enrichment was calculated using the ClusterProfiler package in R. Fold enrichment is calculated by comparing the proportion of genes annotated with a GO term in the cluster relative to the proportion of genes annotated with that GO term in the whole genome. P values are corrected for multiple hypothesis testing using the Benjamini-Hochberg procedure.

### Transmembrane transport gene expression peaks while establishing infection

Cluster 2 encompasses genes whose expression is highest at 1–4 hpi, making this gene set particularly relevant to early infection, where invading *R. parkeri* must evade host responses and form a habitable niche. Notably, this cluster has the highest relative proportion of pseudogenes (61 of 375) and hypothetical genes (177 of 375), suggesting that *R. parkeri* may leverage novel genes during the establishment of infection. Within the remaining 137 genes that had annotated functions, we saw statistically significant enrichment in transmembrane transport-associated genes and genes with efflux transmembrane transport activity (**Fig. 3C**). Though not statistically significant, cluster 4 also contained a modest enrichment in additional transmembrane transport genes, whose temporal pattern also showed the highest expression at the early infection stages (**Fig. 3C**). While the exact nature and regulation of these transporters in *R. parkeri* is unknown, transposon mutagenesis of transporter genes *MC1_RS01115* (cluster 2) and *MC1_RS02755* (cluster 4) appear to attenuate *R. parkeri* by reducing plaque sizes during mammalian cell infections (34), highlighting their potential functional importance.

### Ribosomal, translation, and lipid transport-associated gene expression increases as infection progresses

After the initial ∼ 8 hr lag phase of *R. parkeri* intracellular growth, population expansion ramps up with doubling times every 5–6 hrs (21, 32). Exponential growth calls for an increased production of proteins and rapidly expanding the pool of ribosomes and translation factors is expected to meet those demands (35, 36). Indeed, we observed a statistically significant enrichment of genes encoding ribosome, translation, and RNA binding products in cluster 6 where their expression peaks ≥ 8hpi, concurrent with the bacterium’s exit from lag phase. To accommodate growing protein demands during exponential growth, bacteria must expand their envelope capacity (37). Accordingly, cluster 5, whose genes showed their lowest expression at 1–4 hpi but gradually increased as replication commenced, contained the highest fold enrichment value of any GO term, which was assigned to lipid transport genes. *R. parkeri* genes in this set included homologs of lipopolysaccharide transporters *lptC* (MC1_RS03380) and *lptF* (MC1_RS06065), lipid II transporter *murJ* (MC1_RS04585), and a putative *asmA*-like phospholipid transporter (MC1_RS02440). These data may reflect the temporal transcriptional control of a subset of annotated lipid transport genes that likely serve to assemble and maintain the cell envelope during replication.

### Gene products that potentiate host interactions show diverse transcriptional trends

To complement our systems-level analysis of rickettsial gene networks, we next examined specific genes known to promote intracellular infection. *R. parkeri* employs secreted and surface proteins to interact with and manipulate the intracellular host environment, and several of these proteins are known to function at specific intracellular life cycle stages (**Fig. 4A**). For example, SFG *Rickettsia* execute two temporally and morphologically distinct phases of actin-based motility, either driven by the surface protein RickA to form short tails during the first hour of infection or the autotransporter Sca2 to form long tails later in infection (20). Despite the temporal differences in protein function, the transcriptional trends of *rickA* and *sca2* are remarkably similar. These genes were assigned to cluster 6 and their expression increases from 0–4 hpi and then remains steady, with modest fluctuations (**Fig. 4B**). These data agree with prior work showing that both proteins are expressed steadily throughout infection (20).

**Fig. 4.**
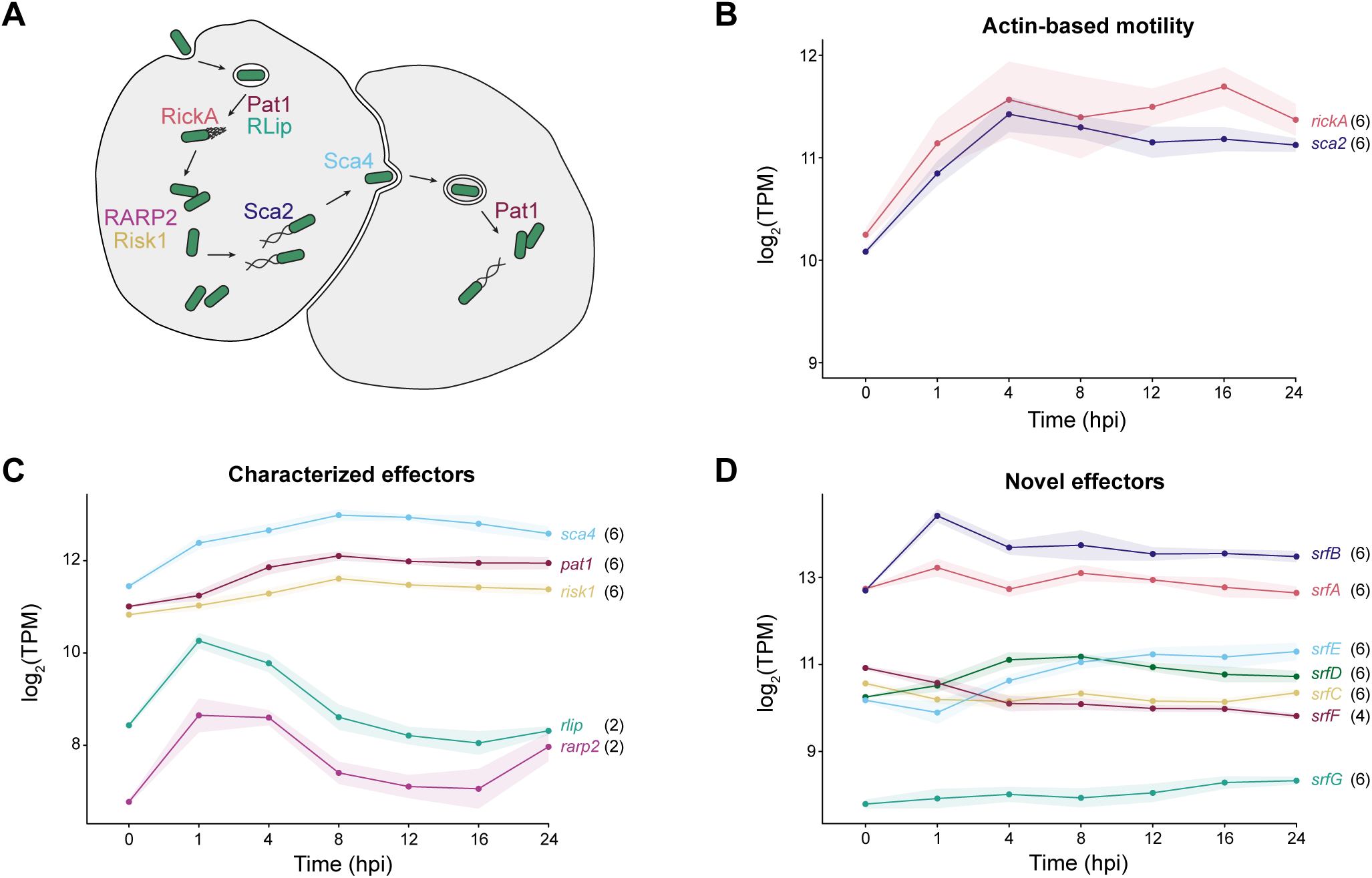
Bacterial gene products that facilitate host cell manipulation follow variable expression patterns. (**A**) Depiction of *R. parkeri* infectious life cycle stages and the bacterial gene products involved at each step. (**B**) Expression trends of bacterial genes involved in actin-based motility, (**C**) characterized bacterial secreted effectors, and (**D**) novel secreted effectors of unknown function. TPM values are averaged across 6–8 biological replicates per time point and shaded region represents standard deviation. Cluster assignments are denoted in black next to each gene name.

Cluster 6, whose genes are defined by increased expression over time, also showed enrichment of genes encoding secreted effectors (GO:0052048) (**Fig. 3C**), leading us to further investigate the expression pattern associated with each effector. We first investigated experimentally validated secreted effectors in *R. parkeri*, including those with known functions like Pat1 (38), Risk1 (39), Sca4 (40, 41), RLip (42), and RARP2 (43) (**Fig. 4C**). The effector-encoding genes assigned to cluster 6 produce products such as the patatin-like phospholipase A2 enzyme (Pat1), which is required for vacuole escape early and late in infection (38), the phosphatidylinositol 3-kinase effector Risk1, which modulates host intracellular trafficking (39), and the cell-to-cell spread effector Surface cell antigen 4 (Sca4) (**Fig. 4C**). In contrast, the genes encoding RLip and RARP2 were assigned to cluster 2, as their expression peaked early in infection (1 hpi) and decreased after 4 hpi (**Fig. 4C**). This expression pattern is consistent with RLip’s role in early phagosomal escape and coincides with previous temporal qPCR data in endothelial cells (42). RARP-2 is thought to modulate host protein secretion (43) but its temporal behavior has yet to be determined. In addition to these effectors of known function, a group of novel *S*ecreted *r*ickettsia *f*actors (SrfA-E and SrfG) are also encoded by genes assigned to cluster 6 (**Fig. 4D**), while SrfF was associated with cluster 4 (**Fig. 4D**). These novel secreted effectors have little to no known functions (23), but their expression patterns may indicate at which stage of infection they are secreted into the host cell.

### Fracturing the T4SS operon yields distinct temporal signatures

Because temporal RNA-seq improves operon prediction and uncovers complex transcriptional relationships, we next examined the operon structures of genes encoding an anomalous secretion system. *Rickettsia* spp. encode a complex type IV secretion system, which is thought to translocate a subset of effectors into the host. Unlike the canonical *Agrobacterium tumefaciens vir* T4SS, which is built from 12 core components and largely encoded on a single operon, the Rickettsiales vir homolog (*rvh*) T4SS has undergone massive gene family expansion and diversification to include duplicated (*rvhB4-I–II*, *rvhB8-I–II*, and *rvhB9-I–II*) and quintupled genes (*rvhB6a*-*e*) (44). Furthermore, the *rvh* genes are dispersed throughout the rickettsial genome, with most of the subunits encoded within two separate genomic subsets, in addition to several scattered genes (**Fig. 5A**). Prior analyses of their operonic structures in related *Rickettsia* spp. predicted divergent transcript boundaries in different species, highlighting a need to define the operon structure in *R. parkeri*. Therefore, we assessed the T4SS operonic structures using Rockhopper (45), which leverages both genome sequence information and expression data to predict operon structures (**Table S5**).

**Fig. 5.**
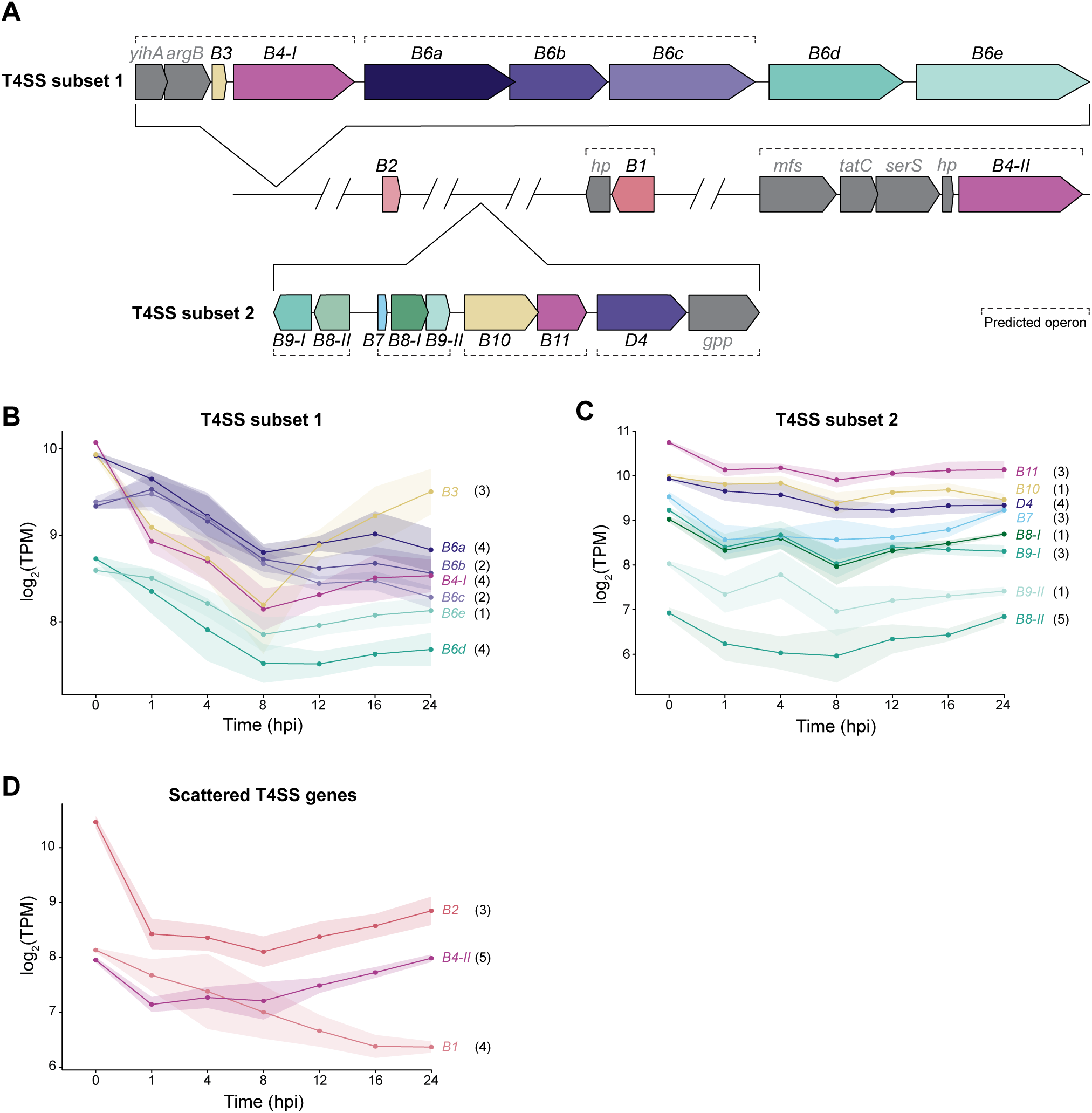
T4SS subunits display diverse expression patterns, regardless of genome position. (**A**) Genome diagram of *R. parkeri* showing the relative locations of T4SS subset 1, subset 2, and scattered genes. Predicted operon structure is annotated by dashed lines. Genes in grey are not part of the T4SS. *hp* = hypothetical protein; *mfs* = MFS-superfamily transport protein; *gpp* = Ppx/GppA phosphatase family protein (**B**) Temporal expression trends of T4SS genes located in T4SS subset 1, (**C**) T4SS subset 2, and (**D**) for the scattered T4SS genes. TPM values are averaged across 6-8 biological replicates per time point and shaded region represents standard deviation. Cluster assignments are denoted in black next to each gene name.

The first genomic subset of T4SS genes contains genes that encode the inner membrane and cytosolic components RvhB3, RvhB4-I, and all 5 RvhB6 paralogs. Within this subset *rvhB3* and *rvhB4-I* are predicted to be in an operon with the non-T4SS genes *yihA* (a ribosome biogenesis factor) and *argB* (an acetylglutamate kinase), followed by the next operon containing *rvhB6a-c* (**Fig. 5A** and **Table S5**). Correspondingly, *rvhB6a-c* have similar expression levels and are expressed at higher levels than *rvhB6d*-*e* (**Fig. 5B**). In contrast, *rvhB3’s* expression diverges from the other members of its operon ≥ 12hpi, indicating a more complex relationship. The second T4SS subset contains genes encoding the structural components RvhB7-10 and the ATPases RvhB11 and RvhD4, and are predicted to be encoded in 4 different operons (**Fig. 5A**). Intriguingly, we see tightly correlated expression between *rvhB8-I* and *rvhB9-I* (**Fig. 5C**), even though these genes are in opposite orientations and separated by 2 other genes, *rvhB8-II* and *rvhB7*. The remaining T4SS components, *rvhB1*, *rvhB2*, and *rvhB4-II*, are all in isolated genomic regions, each with variable incorporation into predicted operons with non-T4SS genes (**Fig. 5A**) and diverse expression trends (**Fig 5D**). Our data collectively show that transcriptional regulation of the *Rickettsia* T4SS differs dramatically from the simpler *Agrobacterium* T4SS, which likely correlates with the structural complexity inherent within the expanded *rvh* system.

### Antisense RNAs show diverse expression patterns and relationships to corresponding mRNA

The relatively low number of transcriptional regulators predicted to be encoded in the rickettsial genome suggests alternative forms of transcriptional regulation. Post-transcriptional regulation via noncoding RNAs allows bacteria to rapidly tune gene expression. Putative regulatory RNAs have been predicted in some *Rickettsia* spp. (6, 7), but the repertoire of regulatory small RNAs was unknown for *R. parkeri*. Regulatory RNA molecules can be difficult to define, due to technical and biological noise in the bacterial transcriptome (46, 47). *Rickettsia* spp. also exhibit poor transcription termination (48), which obscures transcript boundaries. Therefore, we focused on *cis*-encoded small RNAs (i.e. antisense RNAs, asRNA), which are transcribed from the opposite strand of their gene target (49). We prioritized the relationships between asRNA molecules and their complementary mRNA molecules, although asRNA can act as a trans-encoded regulator of distant genes (50–52). We applied a more stringent cutoff for novel asRNA molecules than previous reports (53), requiring at least 50 sequential nucleotides with an average read depth of 50 on both strands, and an asRNA:mRNA ratio of > 0.5 across that >50 nucleotide region. This resulted in 310 genes (20.7% of all coding genes) that are associated with putative asRNA transcripts (**Table S6**), consistent with other Gram-negative bacteria where 6-47% of their genes are associated with asRNAs (50).

asRNAs have diverse and complex regulatory effects on their corresponding mRNA, such as stabilizing mRNA molecules (54), targeting mRNAs for degradation, or blocking translation (50). Strong correlation between mRNA and asRNA expression levels is often indicative of a regulatory relationship between the two but it is difficult to link the direction of correlation to the mode of regulation, as positive correlation predominates regardless of mechanism (55). To investigate our putative asRNAs, we calculated the Pearson correlation coefficient for each sense and antisense pair over time (**Fig. 6A**). Most asRNAs (162 genes, 52.26%) were positively correlated with their corresponding mRNA, while a subset (39 genes, 12.58%) was strongly negatively correlated, and the remaining asRNAs (109 genes, 35.16%) lacked strong temporal correlation with their corresponding mRNA. We identified several genes involved in known, temporally regulated processes, including the cell division genes *zapE* (**Fig. 6B**) and *ftsI* (**Fig. 6C**), the DNA polymerase I gene *polA* (**Fig. 6D**), the DNA repair gene *recG* (**Fig. 6E**), and the lysine methyltransferase gene *pkmt2* (**Fig. 6F**), which shields the bacterium from host autophagy (56). These genes showed strong positive or negative correlation between their mRNA and asRNA levels, but further work is necessary to uncover the mechanism driving their interactions.

**Fig. 6.**
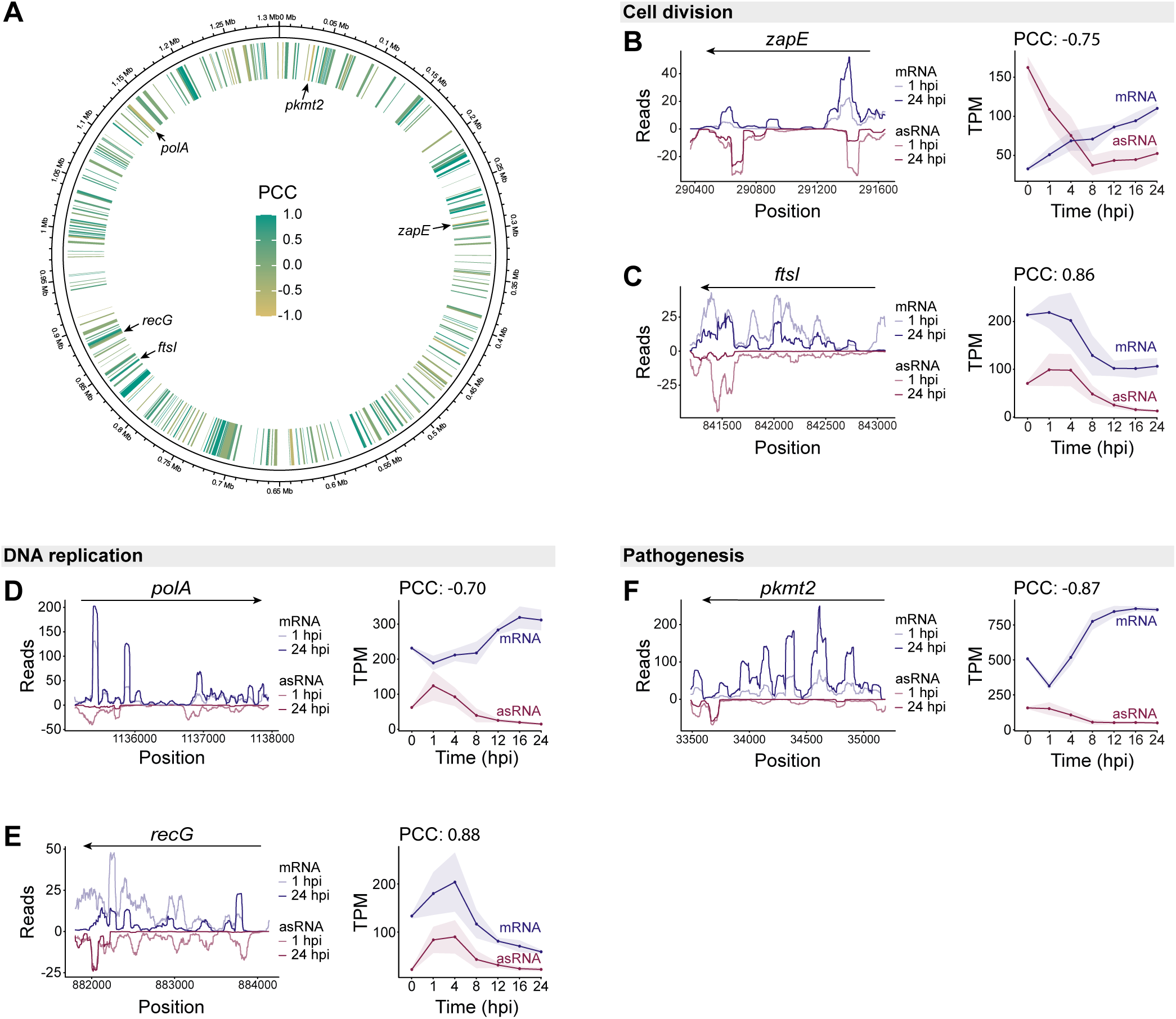
Putative antisense RNA transcripts show variable expression correlation with their cognate mRNA transcripts. (**A**) Genome diagram depicts the positions of 310 putative asRNA transcripts. Transcripts are colored by Pearson correlation coefficient between asRNA and mRNA expression. Average read depth per base pair and TPM over time for the genes *zapE* (**B**), *ftsI* (**C**), *polA* (**D**), *recG* (**E**), and *pkmt2* (**F**). Read depth plots were scaled for each sample by a factor of (1e6/total aligned reads) and averaged across all 1 hpi or 24 hpi replicates. Antisense TPM calculations used the length of the gene’s coding region, because asRNA boundaries were difficult to define.

## DISCUSSION

Prior work profiling the transcriptome of intracellular *Rickettsia* spp. has relied on deep sequencing, long infection timescales, and high bacterial burden. While such conditions are informative, they may obscure nuances in bacterial transcription that occur with physiologically relevant infectious burdens. To address this issue, we deployed PatH-Cap across early and late *R. parkeri in vitro* infection to enrich bacterial transcripts. We achieved higher read depth across the genome with lower bacterial burden than previous works, allowing us to profile infections with as few as a single bacterium per host cell. Using this method, we have generated a complete and robust gene expression dataset in *R. parkeri* that captures early stages of infection and reveals expression changes as infection progresses. With these data we were able to cluster genes based on temporal trends, improve operon predictions for *R. parkeri*, and identify putative asRNAs. We hope this dataset will function as a resource for future studies exploring the transcriptional profile of specific genes or global shifts upon genetic or environmental perturbations.

Our approach allowed us to highlight cohorts of genes that may play roles at specific infection stages. For example, genes enriched in cluster 2 showed the highest expression at 1–4 hpi, which has been historically difficult to profile using standard RNA preparation methods. This cluster is dominated by a set of efflux pumps, which can transport antibiotics out of the bacterial cytoplasm to confer resistance (57), or in some cases promote bacterial growth and virulence (58). While their exact functions remain a mystery, SFG rickettsiae encode significantly more transmembrane transport genes than other rickettsial groups (3), suggesting they may serve a SFG-specific purpose. To further investigate the regulation of genes within this cluster, we attempted 1) motif identification in regions upstream of clustered genes (59) and 2) hub gene analysis using the WGCNA results to identify highly interconnected genes that often play a role in gene regulation (60–62). While we identified many modestly-associated motifs, most genes within the cluster were far too interconnected to reveal potential hub genes. Similar limitations were observed with the other clusters. Future work investigating co-expression across more conditions may be necessary to unravel these regulatory networks throughout the *R. parkeri* infectious life cycle.

In addition to highlighting gene networks of interest, transcriptional profiling has the potential to improve genetic approaches in the rickettsial field. For example, many existing genetic tools for *Rickettsia* spp. use only a small number of promoters and expanding this repertoire to incorporate promoters with different strengths or timing will enable more precise dosage- or stage-specific expression of genes of interest. We also note that the expression trends for common rickettsial housekeeping genes used in qPCR experiments (*adr1*, *ompB*, *gltA*, *rpoD*) (16, 42, 44) require re-evaluation. Both *adr1* and *ompB* expression more than doubled over the course of infection, *gltA* expression modestly increased (log_2_(FC) = 0.58), and *rpoD* was stable across time (log_2_(FC) = 0.25), highlighting the importance selecting the correct controls. Moreover, the identification of species-specific asRNAs should motivate studies untangling the regulatory relationships between mRNA and asRNA. asRNA systems have been manipulated in other bacterial species for targeted gene knockdown (63–65) and prior work in the Rickettsiaceae family member *Orientia tsutsugamushi* has shown that high levels of asRNA correlate with reduced protein levels (8). Thus, these naturally occurring asRNAs could serve as the basis for genetic tools in *Rickettsia* spp.

Despite its streamlined genome, *Rickettsia* spp. encode many pseudogenes and hypothetical proteins of unknown function, limiting the deeper functional predictions possible from transcriptomics in well-studied systems. Although our analysis uncovered transcriptional trends for these enigmatic genes and establishes that many of these loci are expressed during infection, proteomic analysis is required to determine whether these genes encode functional proteins.

Our dataset improves operon predictions by supplying information on expression from intergenic regions and co-expression changes between neighboring genes. However, these predictions do not supersede experiments like long-read sequencing to validate operon structures. Additionally, high-throughput methods like end-enriched RNA-seq (66) could profile transcription start and stop sites across the genome and potentially unravel the complex expression patterns we observed within operons, including those encoding the T4SS components.

Our successful application of PatH-Cap to profile *R. parkeri* gene expression enables future work exploring acute rickettsial gene expression changes, including across diverse host- and vector-derived cell types and tissue samples. Alongside targeted studies of genes of interest, PatH-Cap provides a complementary approach to illuminate how rickettsiae successfully navigate distinct environments and cause disease. Simultaneously sequencing the host or vector transcriptome alongside PatH-Cap RNA-seq, similar to what has been done with *M. tuberculosis* (18), could also highlight points of crosstalk between these systems. Together, these analyses should address how bacterial and host/vector transcription changes at the population level, which should inform future work dissecting the heterogeneity of these transcriptional changes using single-cell RNA-seq techniques rapidly emerging in the field.

## METHODS

### Mammalian cell culture

A549 human lung epithelial and Vero monkey kidney epithelial cell lines were obtained from the University of California, Berkeley Cell Culture Facility (Berkeley, CA). A549 cells were maintained in Dulbecco’s modified Eagle’s medium (DMEM; Gibco 11965118) supplemented with 10% fetal bovine serum (FBS). Vero cells were maintained in DMEM supplemented with 5% FBS. Cell lines were confirmed to be mycoplasma-negative in a MycoAlert PLUS assay (Lonza, LT07-710) performed by the Koch Institute High-Throughput Sciences Facility (Cambridge, MA).

### Propagation of *R. parkeri*

Wild-type *R. parkeri* strain Portsmouth (kindly provided by Chris Paddock) was propagated by infection of Vero cells grown in DMEM supplemented with 2% FBS at 33°C. Bacteria were isolated through mechanical disruption of infected cells as previously described (40) with additional passage through a sterile 2 μm filter (Cytiva 6783-2520). Isolated bacteria were aliquoted and stored at –80 °C. Bacterial titers of frozen stocks were determined by plaque assay (34).

### Infection and RNA preparation

For all assays, A549s were grown in 6-well plates and infected with *R. parkeri* at an MOI of 1. For temporal profiling, the 6–8 biological replicates were collected on the same day to minimize batch effects (67). *R. parkeri* were added to media and spun at 200 x g for 5 minutes to settle bacteria onto the host cell monolayer, then incubated at 33°C until the time point of interest. RNA was collected by aspirating cell culture medium and adding 500 µL of Trizol Reagent (Thermo Fisher Scientific 15596026) to each well. Each replicate used material from 1–2 wells. 0 hpi samples were collected by adding purified *R. parkeri* directly to Trizol Reagent. RNA was extracted using PureLink RNA mini kit (Thermo Fisher Scientific 12183018A) following manufacturer’s Trizol extraction protocol.

### Hybrid capture array design

A custom target enrichment panel was designed for the *R. parkeri* strain Portsmouth NC_017044 genome (April 2017 annotation) in collaboration with Twist Bioscience. Briefly, probes were designed to tile the *R. parkeri* genome end-to-end. The set of probes was checked for matches to rRNA and tRNA transcripts based on the 2017 Los Alamos assembly as well as any match to the human genome (hg38), as checked with salmon v.1.10.0 (68) and bwa mem v0.7.16a-r1181 (69) / bedtools v 2.26.0 (70). A/T nucleotide probe content spanned 45–90% with a mean +/-SD of 67.7%+/- 6.4%, consistent with *R. parkeri*’s genomic AT content of 67.5%. The resulting set of probes, excluding the aforementioned elements, was redesigned and balanced for nucleotide content (available as TE-95904079, Twist Bioscience).

### Library preparation, hybrid capture reaction, and sequencing

Total RNA from infected host cells was used to create stranded, barcoded cDNA libraries using Twist RNA Library Prep Kit (Twist Biosciences 107061) according to manufacturer’s protocol. cDNA libraries from individual biological replicates were generated, normalized by mass, and pooled for capture using the Twist standard hyb and wash kit v2 (Twist Biosciences 105561), our custom *R. parkeri* probe panel (Twist Biosciences), and Twist’s target enrichment standard hybridization v2 protocol. Equal amounts of captured libraries were reamplified, pooled and sequenced as 75 bp paired end reads on an AVITI24 platform (Element Biosciences) at the MIT BioMicro Center (Cambridge, MA).

### Read Mapping and Gene Counts

Reads were mapped to the *Rickettsia parkeri* str. Portsmouth genome (NCBI Reference Sequence: NC_017044.1) using STAR (71). Reads were assigned to genes using featureCounts (72) from the subread module (73) on BioConda (74).

### Analysis of Gene Expression

Genes with fewer than 48 total reads across all replicates and time points were removed. Genes that were not included in the hybrid capture library (rRNAs and tRNAs) were also removed. Depending on downstream analyses, TPMs were calculated or gene counts were normalized and transformed using DESeq2’s rlog function (75). The WGCNA (25) package for R was used to assign gene clusters.

### GO term analysis

Protein coding sequences and a list of pseudogenes were extracted from the GenBank file for *R. parkeri*. We used the InterProScan (33) tool on the Galaxy webserver (76) to assign GO terms to all protein coding genes. Terms were further refined based on literature searches (77). GO term enrichment was calculated using the enricher function in the ClusterProfiler (78) package on R. Reported p values are corrected for multiple hypothesis testing with the Benjamini-Hochberg procedure (79).

### asRNA identification

To identify putative antisense RNAs, we used Samtools (80) to separate first in pair reads that aligned to the forward and reverse strands of the genome, followed by Bedtools (70) from BioConda to generate forward and reverse read depth at the nucleotide level. Putative antisense reads were identified using a custom R script based on the conditions described in the text.

## ACKNOWLEDGEMENTS

We thank Alexei Stortchevoi, Duanduan Ma, and the staff at the MIT BioMicro Center for technical assistance, guidance, and cDNA library preparation. We also thank members of the Lamason lab for help with sample collection and critical reading of the manuscript. This work was supported in part by National Institutes of Health Grants no. T32GM007287 and GM136540 (ATS), and R01AI155489 (RLL), and the Damon Runyon Cancer Research Foundation (JM).

## DATA AVAILABILITY

RNA sequencing data are available through NCBI’s Gene Expression Omnibus accession number GSE314162.

---

**Fig. S1.**
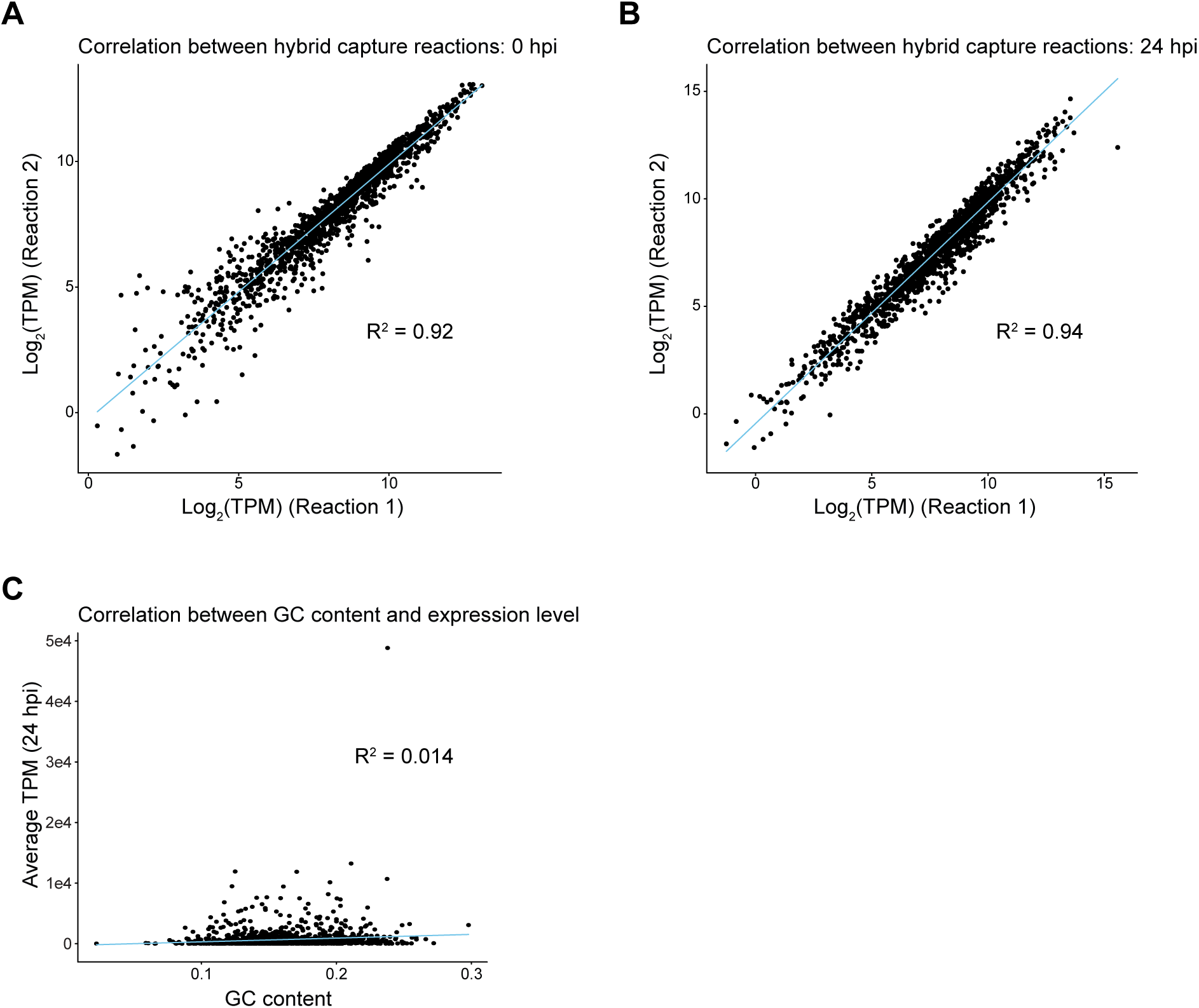
Hybrid capture reactions are reproducible and unaffected by gene GC content. Correlation between hybrid capture reactions of RNA samples collected from biological replicates at 0 hpi (**A**) and 24 hpi (**B**). Each reaction contained 6–8 biological replicates per time point and TPM values were calculated as an average of those replicates. (**C**) Correlation between GC content and gene expression level at 24 hpi.

**Fig. S2.**
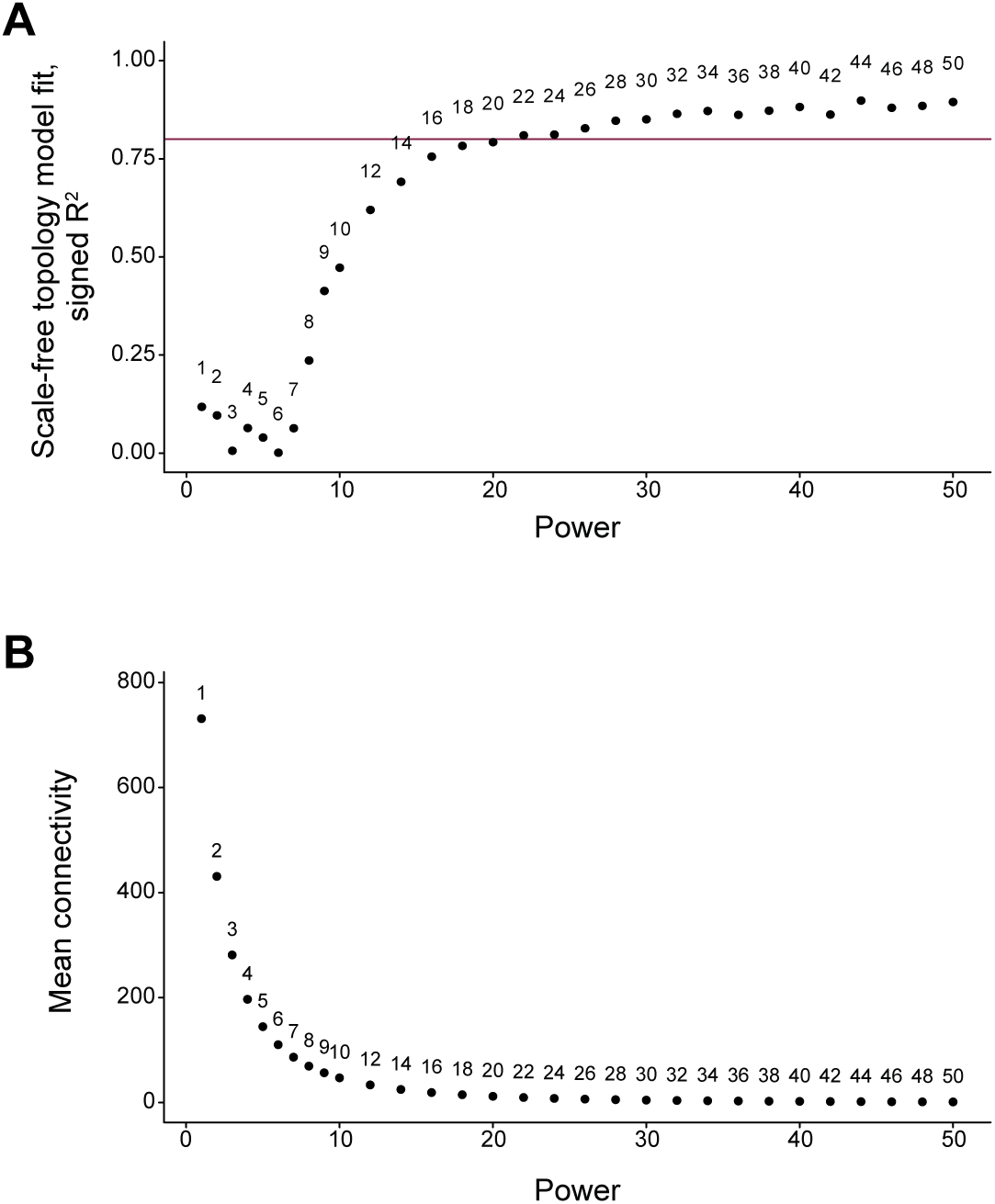
Assessment of soft-thresholding power parameter for WGCNA. (**A**) Varying values of the soft-thresholding power parameter (β) and the corresponding scale-free topology model or (**B**) the corresponding mean connectivity values. β was set to 22, which was the smallest power value to fit the scale-free topology model with R^2^ > 0.8 (red line) that also minimized the mean connectivity value.

**Fig. S3.**
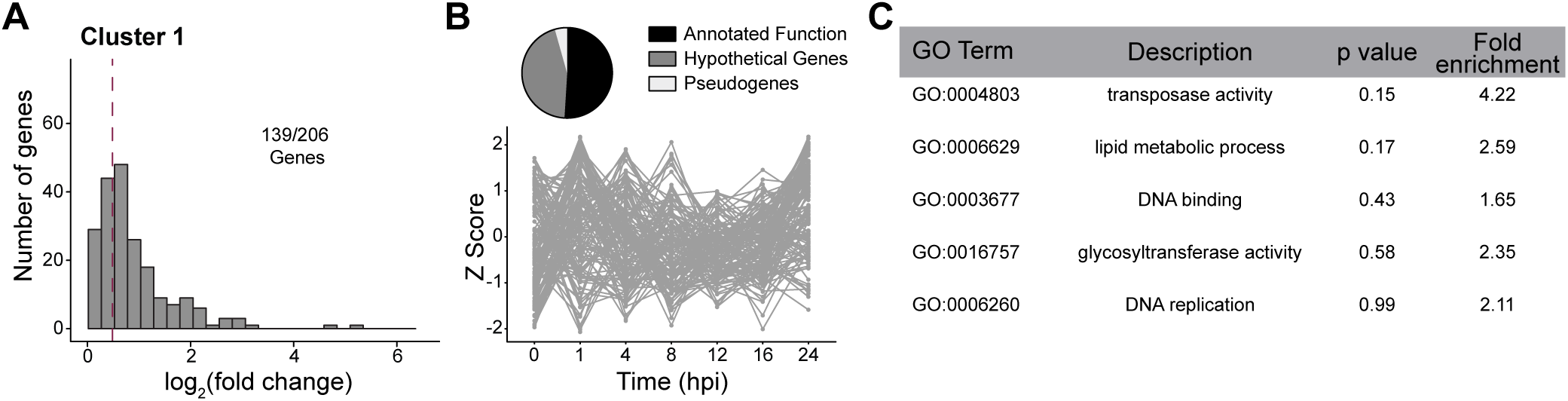
Temporal changes and gene content for cluster 1. (**A**) Histogram log_2_(FC) for genes in each cluster. Maximum log_2_(FC) values were calculated as described for Figure 3. (**B**) Pie charts show the proportion of genes within a cluster annotated as genes of annotated function, hypothetical genes, and pseudogenes. Line graph shows Z score over time of genes in the cluster. (**C**) Top 5 enriched GO terms within each cluster, reported regardless of statistical significance.

**Table S1** DESeq2 analysis of differentially expressed genes comparing 24 and 48 hpi or 24 and 0 hpi. Differentially expressed genes displayed in Fig. 1D are denoted by text color.

**Table S2** Genes with the highest and lowest average expression across all time course samples and their expression in different conditions from previous works.

**Table S3** Cluster and pseudogene, hypothetical, or GO term assignments for all genes in the *R. parkeri* genome included in this work.

**Table S4** GO term enrichment data for all clusters as calculated by ClusterProfiler.

**Table S5** Operon prediction based on Rockhopper analysis. Genes not listed in this table were predicted to not be in an operon.

**Table S6** Correlation, asRNA:mRNA ratio, and associated GO terms for all 310 genes with associated putative asRNA transcripts.

